# inteRelate: flexible and thorough relating of genomic interval datasets through comparative overlap analysis

**DOI:** 10.64898/2026.09.14.751391

**Authors:** Aaron Mamane-Logsdon, Mphatso D. Kalemera, Goedele N. Maertens

**Author notes:** Corresponding authorship.

## Abstract

**Summary:** Testing spatial relationships between genome-mapped features is both a common source of hypothesis generation and an additional layer of supporting evidence for experimental findings. Many computational tools automate the statistical association procedures used to assess overlap between pairs of genomic features. However, none provides a dedicated workflow that directly compares multiple features by assessing their overlap with a separate, common feature, first testing for overall heterogeneity and then identifying which overlap rates differ. Such comparative overlap analysis could directly facilitate comparisons of features within the same class across disease states, cell types, experimental perturbations, and other biological contexts. Here, we describe inteRelate, a software package that uses genomic interval datasets to test spatial relationships between genome-mapped features through comparative overlap analysis. The package functions as an end-to-end pipeline that is highly tunable and thorough in its statistical association procedures. We use experimental data to demonstrate the automation, accuracy, and insight inteRelate provides.

**Availability and implementation:** inteRelate is available at https://github.com/loggy01/interelate and archived at https://zenodo.org/records/21891072. Example uses are available in the online supplement. Additionally, the example datasets and results are available at https://zenodo.org/records/22012767.

## 1 Introduction

Experimental approaches that map a feature to a reference genome commonly produce a dataset of genomic intervals, each interval an instance of that feature defined by its chromosome and start-end coordinates, a representation popularised by the Browser Extensible Data (BED) format (Niu *et al*. 2022) developed at UCSC for the Human Genome

Project (Lander *et al*. 2001) and supported by the UCSC Genome Browser (Kent *et al*. 2002). These datasets can be used to generate hypotheses or to provide additional evidence for experimental findings by relating one to another through spatial association of their genomic intervals, thereby associating features of interest with other features such as genes; cis-regulatory elements and binding sites for trans-acting regulators; epigenomic features such as chromatin accessibility, histone modifications, and DNA methylation; higher-order chromatin features; evolutionarily conserved regions; and genetic variants, disease-associated loci, and more. This utility has motivated several established pipelines that automate and statistically support spatial association of genome-mapped features. Most of these frameworks statistically test overlap or other spatial associations between feature pairs against null distributions derived analytically (Favorov *et al*. 2012, Sheffield and Bock 2016, Layer *et al*. 2018) or through randomisation (Favorov *et al*. 2012, Heger *et al*. 2013, Gel *et al*. 2016). While these frameworks extend beyond independent pairwise association tests to varying degrees, none directly compares multiple features by jointly testing for heterogeneity in their overlap rates with a separate, common feature and then, where heterogeneity is detected, identifying which overlap rates differ. We believe this comparative overlap analysis is important because it could directly facilitate comparisons of features of the same class across disease states, cell or tissue types, developmental stages, time points, experimental treatments or perturbations, and more. Accordingly, comparative overlap analysis would also benefit from a robust, end-to-end statistical pipeline.

To address this gap, we developed inteRelate, a Python package for comparative overlap analysis of genome-mapped features using genomic interval datasets. The package provides a convenient end-to-end pipeline that flexibly processes input datasets, quantifies interval overlap at user-specified genomic distance thresholds, jointly tests multiple datasets for overall heterogeneity in their observed overlap rates with a common dataset, identifies which overlap rates differ where heterogeneity is detected, and collates the overlap rates and statistical results in a unified, human-readable report. Additionally, inteRelate scales to run multiple comparative overlap analyses in a single run. It promotes comprehensive interpretation by combining complementary measures, including overlap rate, statistical significance, and association strength. Its statistical procedures are highly configurable, allowing users to select and fine-tune tests to complement their data’s properties and their experimental design.

## 2 Software description

### 2.1 Implementation

inteRelate is a pip-installable Python package that runs on Linux, macOS, and Windows. Users can run the entire workflow through the “interelate” command, providing a convenient end-to-end pipeline for comparative overlap analysis. However, inteRelate remains flexible through user-tunable overlap-counting and statistical-testing modules. Additionally, its open-source implementation provides substantial support to developers, with extensive documentation of its test suite design that aids collaborative development of the package’s functionality to better suit user needs.

### 2.2 Counting interval overlaps

Consistent with inteRelate’s flexibility, the overlap-counting module accepts genomic interval datasets in any BED format from BED3 to BED12 (Niu *et al*. 2022), including gzip-compressed files. inteRelate requires at least one query dataset to serve as a common feature against which it measures overlap for each reference dataset. There must be at least two reference datasets, and one is advised to be an appropriate control from which overlap enrichment above background levels can be measured. Users may also specify multiple genomic distance thresholds to define overlap. The module treats the reference datasets, query datasets, and genomic distances as finite, indexed families and evaluates every possible combination formed by selecting the next member from each family: first, the *i*-th reference dataset *R*_*i*_; second, the *j*-th query dataset *Q*_*j*_; and third, the *k*-th genomic distance *d*_*k*_. For each combination, using PyRanges1 (Stovner *et al*. 2025), the module counts, for every interval in *R*_*i*_, the number of intervals in *Q*_*j*_ that lie within *d*_*k*_ base pairs, with counts for all supplied values of *d*_*k*_ collated into a separate text report for each fixed pair of *R*_*i*_ and *Q*_*j*_ (Fig. 1, Stage 1), providing transparency in overlap counting.

**Fig. 1.**
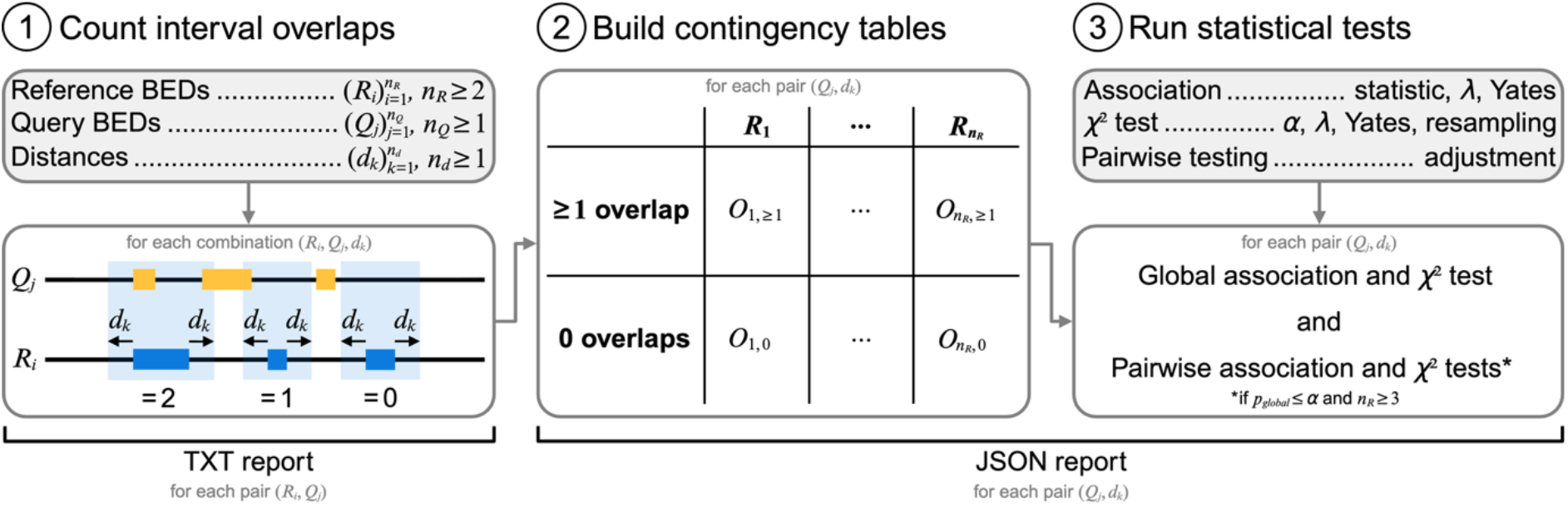
Schematic overview of how genomic interval datasets are related through inteRelate. The constituent boxes outlined in grey represent either user input (grey fill) or internal processes (white fill). Output reports and the processes they are derived from are indicated by square brackets. Grey annotations indicate the scope of the corresponding processes and outputs. The schematic comprises three stages of comparative overlap analysis: (1) count interval overlaps, (2) build contingency tables, and (3) run statistical tests. In the first stage, the overlap-counting module takes three inputs: reference BED datasets, query BED datasets, and genomic distances. Each input is represented as a finite, indexed family, where the index identifies an individual member and the upper index limit denotes the family size. This stage evaluates every combination of one reference dataset *R*_*i*_, one query dataset *Q*_*j*_, and one genomic distance *d*_*k*_. For each combination, every interval in *R*_*i*_ (blue) is extended by *d*_*k*_ at both ends, and the number of intervals in *Q*_*j*_ (yellow) overlapping the resulting region (translucent blue) is counted. For each fixed pair of *R*_*i*_ and *Q*_*j*_, overlap counts for each *d*_*k*_ are recorded in a single report. In the second stage, to build a contingency table for each fixed pair of *Q*_*j*_ and *d*_*k*_, the table-building module uses the overlap counts from each reference dataset to classify its intervals as either having at least one overlap or no overlaps with that *Q*_*j*_ at that *d*_*k*_, determining the observed frequencies *O*_*i*, ≥ 1_ and *O*_*i*, 0_ respectively. Therefore, reference datasets form the columns and overlap statuses form the rows. In the third and final stage, the statistical-testing module receives several input arguments to fine-tune its testing procedures. For the contingency table from each fixed pair of *Q*_*j*_ and *d*_*k*_, an association statistic and a *χ*^2^ test of independence are calculated. If the *χ*^2^ test is statistically significant and at least three reference datasets are provided, the stage repeats for each two-reference subtable with multiple-testing correction. All test results, together with the observed frequencies and reference-specific overlap rates from the previous stage, are compiled into a single report.

### 2.3 Building contingency tables

For each fixed pairing of *Q*_*j*_ and *d*_*k*_, every reference interval can be described by as few as two categorical variables: the reference dataset to which it belongs and its overlap status with *Q*_*j*_ at *d*_*k*_, categorised as at least one overlap or none. Therefore, a contingency table provides the natural structure for testing whether these variables are associated and, equivalently, whether overlap rates differ among the reference datasets. Accordingly, using NumPy (Harris *et al*. 2020), the table-building module constructs one table for each fixed pairing of *Q*_*j*_ and *d*_*k*_, with reference dataset categories as columns and overlap status categories as rows, meaning within the *R*_*i*_ column, there are the observed frequencies for intervals in *R*_*i*_ with at least one overlap, *O*_*i*, ≥ 1_, and no overlap, *O*_*i*, 0_ (Fig. 1, Stage 2). Therefore, the proportion of the column total represented by *O*_*i*, ≥ 1_ gives the overlap rate for *R*_*i*_ with *Q*_*j*_ at *d*_*k*_. Placing the overlap rates for all reference datasets in a common table prepares them for direct statistical comparison in the next module.

### 2.4 Running statistical tests

For the contingency table from each fixed pairing of *Q*_*j*_ and *d*_*k*_, using SciPy (Virtanen *et al*. 2020) and Statsmodels (Seabold and Perktold 2010), the statistical-testing module determines heterogeneity among the observed overlap rates of the references and identifies which differ as follows: calculating an association statistic; performing a Pearson’s *χ*^2^ test of independence; repeating both calculations for every two-reference subtable with multiple testing correction, provided the previous global *p*-value was significant and at least three reference datasets were provided; and reporting all results, including the observed frequencies and overlap rates, in a JSON file (Fig. 1, Stage 3). First, we selected Pearson’s *χ*^2^ test of independence because calculating *p*-values from the asymptotic *χ*^2^ null distribution of Pearson’s test statistic is generally more computationally tractable than exact inference as contingency table dimensions increase (Agresti 1992), and thus scales better as the number of reference datasets increases. However, for sparse contingency tables, this asymptotic distribution may not accurately approximate the null sampling distribution of Pearson’s statistic (Agresti and Yang 1987). Therefore, users can opt to retain Pearson’s *χ*^2^ statistic while replacing the asymptotic *p*-value calculation with an estimate based on either permutation or Monte Carlo resampling under the null hypothesis of independence. Alternatively, for tables representing only two reference datasets, users can apply Yates’ continuity correction to the asymptotic test to achieve a more conservative *p*-value (Yates 1934). Additionally, for asymptotic tests, users may specify *λ* to replace Pearson’s *χ*^2^ statistic with another member of the Cressie-Read power-divergence family, thereby changing how discrepancies between observed and expected frequencies contribute to the statistic (Cressie and Read 1984). Second, statistical significance does not quantify the strength of association between reference dataset identity and overlap status. Therefore, the module complements each *χ*^2^ test with a user-selected association statistic: Cramér’s *V*, Tschuprow’s *T* or Pearson’s contingency coefficient. Where applicable, any selected Yates’ correction or *λ* setting is also applied here. Third, regarding pairwise testing on two-reference subtables, users may select from a wide range of *p*-value adjustment methods to correct for multiple testing. Finally, the JSON report compiled from the contingency table maximises readability by separating results into three distinct, intuitive sections: first, “overlap_result”, which contains the observed frequencies and overlap rates; second, “global_testing_result”, which contains the association statistic and *χ*^2^ test from the whole contingency table; and third, “pairwise_testing_result”, which is the same as the previous section but for each two-reference subtable instead, with the additional adjusted *p*-values. Therefore, a single report shows whether overlap rates are heterogeneous across the reference set, identifies which overlap rates differ significantly, and quantifies the strengths of whole and subtable associations; the reference-specific overlap rates provide the context needed to interpret these results. We provide an example of the full JSON schema (Fig. S1).

## 3 Results

To demonstrate inteRelate’s ability to automate accurate comparative overlap analysis of genomic interval datasets, we present two examples based on the genomic integration site (IS) patterns of the retroviruses human immunodeficiency virus type 1 (HIV-1) and human T cell lymphotropic virus type 1 (HTLV-1) (Supplementary note S1). First, compared with matched random control (MRC) ISs, inteRelate reproduced the known associations of both viruses’ ISs with H3K36me3 marks and suggested HIV-1’s association was more significant and stronger (Fig. S2A; Table S1 i), consistent with the direct and indirect roles that H3K36me3 marks have in IS selection for HIV-1 (Sapp *et al*. 2022, Hope *et al*. 2026) and HTLV-1 (Melamed *et al*. 2022) respectively. Second, compared with MRC ISs, inteRelate reproduced the known associations of both viruses’ ISs with the binding sites of the Ser2-phosphorylated, elongating form of RNA polymerase II (POLR2AphosphoS2) and suggested that both viruses’ associations were equal (Fig. S2B; Table S1 ii), consistent with the indirect role of POLR2AphosphoS2 in IS selection for both viruses (Melamed *et al*. 2022, Singh *et al*. 2022).

## 4 Conclusion

inteRelate is the first computational tool to support comparative overlap analysis of genomic interval datasets by providing an intuitive, end-to-end pipeline that streamlines statistical testing for heterogeneity in overlap rates among a group of genome-mapped features with a separate feature at varying genomic distances, and encourages thorough analysis by combining complementary measures of association. Additionally, inteRelate can effectively automate the task by performing multiple comparative overlap analyses in a single run. Further, the flexibility of inteRelate is demonstrated by the diverse BED formats it accepts and the configurability of its overlap-counting and statistical-testing modules. Future development of inteRelate may include expanding its statistical procedures and comparing new genomic spatial associations. Given its broad applicability, we anticipate that inteRelate will support comparative overlap analyses of genome-mapped features across diverse areas of biological research.

## Acknowledgements

We thank the members of the Maertens laboratory for their helpful discussions over inteRelate.

## Author contributions

Conceptualisation: A.M.-L., M.D.K., G.N.M. Data curation: A.M.-L. Formal analysis: A.M.-L. Funding acquisition: A.M.-L., G.N.M. Investigation: A.M.-L. Resources: A.M.-L., M.D.K., G.N.M. Methodology: A.M.-L., M.D.K., G.N.M. Project administration: A.M.-L., G.N.M. Software: A.M.-L. Supervision: A.M.-L., G.N.M. Validation: A.M.-L. Visualisation: A.M.-L. Writing – original draft: A.M.-L. Writing – review & editing: all authors. All authors read and approved the final manuscript.

## Supplementary material

Supplementary material is available online.

## Conflict of interests

None declared.

## Funding

This work was supported by funding from Imperial College London President’s PhD scholarships (A.M.-L) and a grant from Blood Cancer UK (22002 to G.N.M.).

## Data availability

inteRelate is available at https://github.com/loggy01/interelate/ and archived at https://zenodo.org/records/21891072. Additionally, the example datasets and results reported here are available at https://zenodo.org/records/22012767.

## Notes

https://zenodo.org/records/22012767

### Competing Interest Statement

The authors have declared no competing interest.

### Summary of Updates

Updated corresponding author to Aaron Mamane-Logsdon (please could Goedele N. Maertens be added as an additional corresponding author)

