## Supplementary data for "inteRelate: flexible and thorough relating of genomic interval datasets through comparative overlap analysis"

```
{
  "overlap_result": {
    "Observed_frequencies": {
      "overlap": [
        ["ref1", ...],
        ["ref2", ...],
        ["ref3", ...]
      ],
      "no_overlap": [
        ["ref1", ...],
        ["ref2", ...],
        ["ref3", ...]
      ]
    },
    "overlap_rate": [
      ["ref1", ...],
      ["ref2", ...],
      ["ref3", ...]
    ]
  },

  "global_testing_result": {
    "expected_frequencies": {
      "overlap": [
        ["ref1", ...],
        ["ref2", ...],
        ["ref3", ...]
      ],
      "no_overlap": [
        ["ref1", ...],
        ["ref2", ...],
        ["ref3", ...]
      ]
    },
    "chi2_statistic": ...,
    "dof": ...,
    "p_value": ...,
    "reject_null": ...,
    "association_statistic": ...
  },

  "pairwise_testing_result": {
    "expected_frequencies": {
      "overlap": [
        [["ref1", ...], ["ref2", ...]],
        [["ref1", ...], ["ref3", ...]],
        [["ref2", ...], ["ref3", ...]]
      ],
      "no_overlap": [
        [["ref1", ...], ["ref2", ...]],
        [["ref1", ...], ["ref3", ...]],
        [["ref2", ...], ["ref3", ...]]
      ]
    },
    "chi2_statistic": [
      [["ref1", "ref2"], ...],
      [["ref1", "ref3"], ...],
      [["ref2", "ref3"], ...]
    ],
    "p_value": [
      [["ref1", "ref2"], ...],
      [["ref1", "ref3"], ...],
      [["ref2", "ref3"], ...]
    ],
    "adjusted_p_value": [
      [["ref1", "ref2"], ...],
      [["ref1", "ref3"], ...],
      [["ref2", "ref3"], ...]
    ],
    "reject_null": [
      [["ref1", "ref2"], ...],
      [["ref1", "ref3"], ...],
      [["ref2", "ref3"], ...]
    ],
    "association_statistic": [
      [["ref1", "ref2"], ...],
      [["ref1", "ref3"], ...],
      [["ref2", "ref3"], ...]
    ]
  }
}
```


**Fig. S1** Screenshot of a full JSON report generated by inteRelate for reference datasets “ref1.bed”, “ref2.bed”, and “ref3.bed” with query dataset $Q_j$ at genomic distance $d_k$. Data has been replaced with ellipses.

**Supplementary note S1**

We aimed to use inteRelate to accurately and simultaneously conduct four comparative overlap analyses by comparing the overlap rates of human immunodeficiency virus type 1 (HIV-1) in vitro, human T-cell lymphotropic virus type 1 (HTLV-1) in vitro, and matched random control (MRC) integration sites (ISs) with (i) H3K36me3 sites, (ii) H3K36me3 shuffled sites, (iii) Ser2-phosphorylated, elongating form of RNA polymerase II (POLR2AphosphoS2) sites, and (iv) POLR2AphosphoS2 shuffled sites. To do this, we first retrieved IS datasets from the same study (Melamed *et al.* 2022) and converted them to BED format (Niu *et al.* 2022), removing any repeated genomic intervals in the process. Next, using the ENCODE portal (Kagda *et al.* 2025), we retrieved BED files for H3K36me3 (accession: ENCFF432EMI) and POLR2AphosphoS2 (accession: ENCFF847DXY), and, to match the IS datasets, reduced each genomic interval to 1 bp centred on its reported peak and removed repeated intervals. Additionally, to serve as negative controls, we generated versions of the H3K36me3 and POLR2AphosphoS2 BED files, randomised by genomic position, using the shuffle function from BEDTools (Quinlan and Hall 2010), with hg38 assembly chromosome sizes–restricted to chromosomes represented in the regular BED file–from the UCSC Genome Browser (Casper *et al.* 2026), preventing overlapping intervals, and using a random seed of 1 for reproducibility. Finally, we performed a single inteRelate run using the HIV-1, HTLV-1, and MRC BED files as references; the regular and shuffled versions of the H3K36me3 and POLR2AphosphoS2 BED files as queries; and a genomic distance of 500 bp for overlap counting, which we chose because the genomic intervals in the original H3K36me3 and POLR2AphosphoS2 had a median size of ~500 bp. All other settings were default.

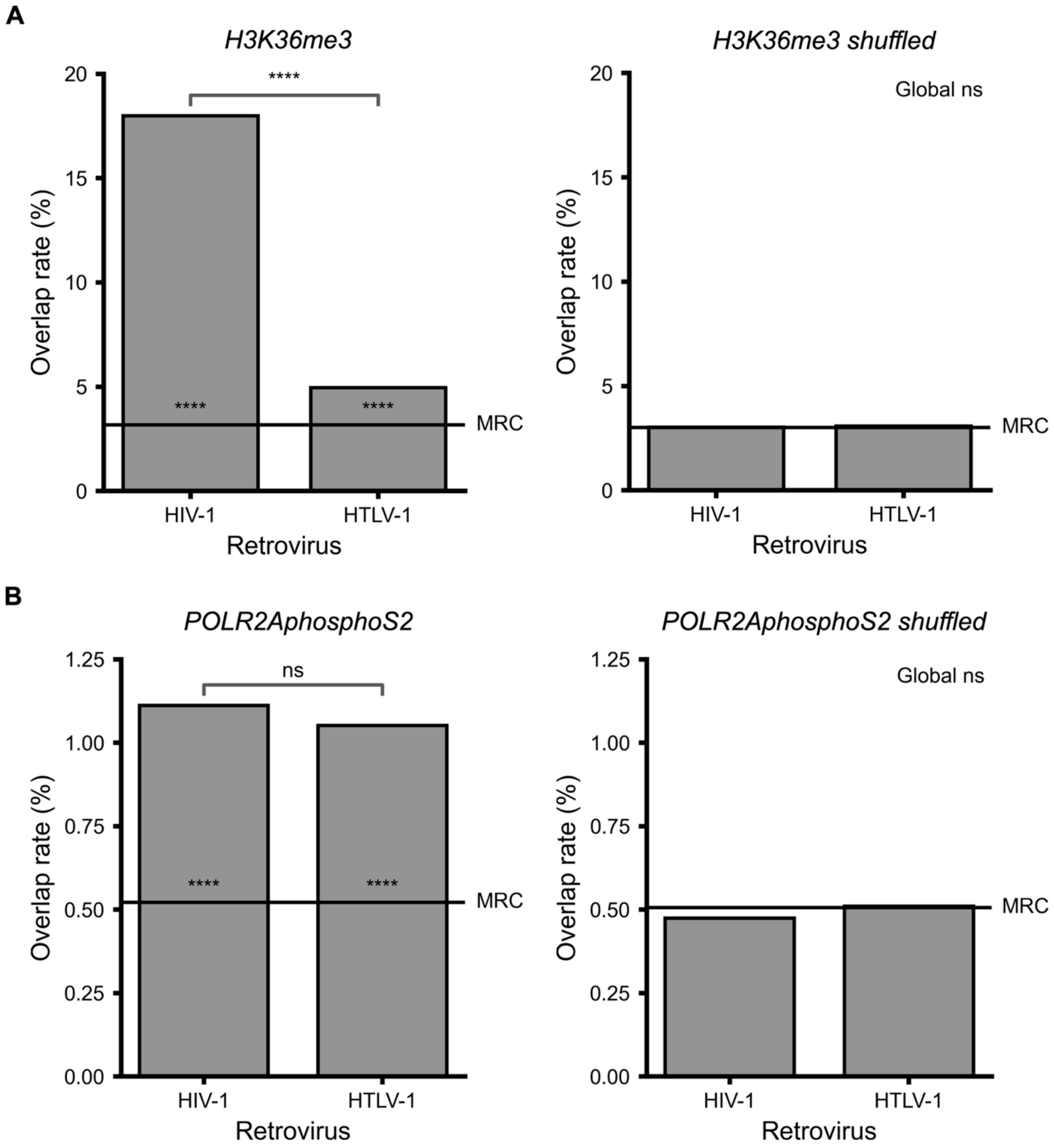


**Fig. S2** Comparisons of how HIV-1 and HTLV-1 ISs relate to H3K36me3 and POLR2AphosphoS2 sites. Overlap rates of HIV-1 and HTLV-1 ISs with **(A)** H3K36me3 sites and **(B)** POLR2AphosphoS2 sites. Data were generated from a single interRelate run using HIV-1, HTLV-1, and MRC BED files as references; H3K36me3 and POLR2AphosphoS2 (regular and shuffled) BED files as queries; and a genomic distance of 500 bp. Overlap rates for HIV-1 and HTLV-1 ISs are shown as bars, whereas the MRC overlap rates are shown as horizontal lines. For plots with a non-significant global Pearson's $\chi^2$ test of independence, "Global ns" is reported in the top-right of the respective plot. Otherwise, follow-up pairwise $\chi^2$ test results are shown as Holm-Šídák-adjusted $p$-values ($q$-values): $q > 0.05$, ns; $q \leq 0.05$, *; $q \leq 0.01$, **; $q \leq 0.001$, ***; $q \leq 0.0001$, ****. Brackets indicate $q$-values between HIV-1 and HTLV-1, whereas $q$-values between each virus and the MRC are shown next to the MRC line.

**(i) H3K36me3**

| Reference one | Reference two | Association statistic |
|---|---|---|
| HIV-1 | HTLV-1 | 0.200 |
| HIV-1 | MRC | 0.254 |
| HTLV-1 | MRC | 0.041 |

**(ii) POLR2AphosphoS2**

| Reference one | Reference two | Association statistic |
|---|---|---|
| HIV-1 | HTLV-1 | 0.002 |
| HIV-1 | MRC | 0.033 |
| HTLV-1 | MRC | 0.026 |

**Table. S1** Pairwise Cramer's $V$ association statistics for HIV-1, HTLV-1, and MRC ISs, using **(i)** H3K36me3 sites and **(ii)** POLR2AphosphoS2 sites as overlap queries. Data were generated from a single interRelate run using HIV-1, HTLV-1, and MRC BED files as references; H3K36me3 and POLR2AphosphoS2 BED files as queries; and a genomic distance of 500 bp.
